# HAT: Haplotype Assembly Tool using short and error-prone long reads

**DOI:** 10.1101/2022.07.20.500775

**Authors:** Ramin Shirali Hossein Zade, Aysun Urhan, Alvaro Assis de Souza, Akash Singh, Thomas Abeel

## Abstract

**Motivation:** Haplotypes are the set of alleles cooccurring on a single chromosome and inherited together to the next generation. Because a monoploid reference genome loses this co-occurrence information, its usability is limited to associate phenotypes with allelic combinations of genotypes. Therefore, methods to reconstruct the complete haplotypes from DNA sequencing data are crucial. Recently, several attempts have been made at haplotype reconstructions, but significant limitations remain. High-quality continuous haplotypes cannot be created reliably, particularly when there are few differences between the homologous chromosomes.

**Results:** Here, we introduce HAT, a haplotype assembly tool that exploits short and long reads along with a reference genome to reconstruct haplotypes. HAT tries to take advantage of the accuracy of short reads and the length of the long reads to reconstruct haplotypes. We tested HAT on the aneuploid yeast strain *Saccharomyces pastorianus* CBS1483 and multiple simulated polyploid data sets of the same strain, showing that it outperforms existing tools.

**Availability:** https://github.com/AbeelLab/hat/

## 1 Introduction

Most eukaryotes have more than one copy of each chromosome, and some species have more than two homologous copies of each chromosome (i.e., polyploids), which is common in plants [9]. In genetics, a haplotype is the combination of individual alleles (one allele of each gene) located on the same chromosome. Because these alleles are located on the same chromosome, they are passed on together to the next generation [**?**]. Haplotype assembly, or reconstruction refers to the task of reassembling each individual haplotype. The need to reconstruct haplotypes arises from the inability of current DNA sequencing technologies, such as next-generation (NGS) and third-generation (TGS) sequencing, to read a chromosome’s sequence from beginning to end. These technologies instead sequence shorter fragments called reads. In addition, chromosome separation before sequencing requires complicated and expensive lab work that is not feasible for most studies. Therefore, it is more common to sequence chromosomes together, and then use computational methods to separate the reads and reconstruct the haplotypes. A monoploid reference genome consists of a mosaic structure of haplotypes with allelic combinations that do not co-occur within any haplotype. Additionally, some of the alleles found in the haplotypes are absent in the monoploid reference. In contrast, with a haplotype-resolved reference, we can understand genetic variation and link phenotypic traits with the associated alleles in the haplotype better.

It is significantly more challenging to reconstruct haplotypes for polyploid genomes than for diploid genomes. If one of the haplotypes of a diploid genome has been phased (i.e., the said haplotype has been inferred), it is trivial to determine the alleles of the other haplotype based on this. On the other hand, in polyploid genomes, other haplotypes may have the same or different alleles [2]. Hence, the phasing of one haplotype does not clearly indicate what alleles are present in other haplotypes.

Recognizing the wide-spread use of NGS and TGS, it is imperative to develop algorithms for haplotype reconstruction from sequencing reads to facilitate various research applications. In the past years, few tools have tackled this problem. nPhase [1] and Whatshap [11] are among some examples of recently developed tools.

This study presents HAT, a haplotype assembly tool that combines short reads and error-prone long reads along with a reference genome to reconstruct haplotypes. Similar to Ranbow [7], HAT first creates seeds from short reads, but then it expands the seeds with long reads. We benchmark HAT against Whatshap and nPhase because both use long reads to phase haplotypes. Using simulated and real yeast genome data, we demonstrate that HAT outperforms both Whatshap polyphase and nPhase in terms of contiguity and the accuracy of phased alleles.

## 2 Methods

### 2.1 Data

Using both simulated and real data is essential to test HAT properly. Simulated data provides the ground truth of haplotypes to evaluate phasing accuracy and the real data validates HAT’s performance.

#### 2.1.1 Simulated data

We use Haplogenerator [8] to simulate haplotypes from the base genome – chromosome ScII of *Saccharomyces pastorianus* CBS1483 with accession number ASM1102231v1 (ChrSc2) [10]. The ground truth is the simulated haplotypes, and ChrSc2 base sequence is the reference. Next, we simulate reads with 20x coverage per haplotype, similar to the simulation design used in previous studies including nPhase. We use Badread [14] version 0.2.0 and and ART [5] version 2.5.8 to simulate reads similar to Oxford Nanopore Technology (ONT) and to Illumina’s HiSeq 2500, respectively. Badread is used with default parameters, ART parameters are available in 7. In total, 6 datasets are generated for ploidy levels 3,4, and 5, with low and high heterozygosity. For the low heterozygosity datasets, we set the parameters of Haplogenerator to produce the same number of SNPs/Insertions/Deletions as the chromosomes ScII, SeIII-ScIII, and ScVIII of CBS1483 which are triploid, tetraploid, and pentaploid respectively. Because chromosomes SeIII-ScIII and ScVIII are smaller than ScII, the chromosome we use for the simulations, we multiply the number of SNPs/Insertions/Deletions by the ratio of genome sizes. For the high heterozygosity datasets, we fit a lognormal distribution on the distances between consecutive SNPs/Insertions/Deletions of chromosome ScII, SeIII-ScIII, and ScVIII and use the parameters on Haplogenerator. The parameter settings for Haplogenerator are in 6.

Then, the short reads are mapped to the monoploid reference genome using BWA-MEM (Li, 2013) with default parameters. We obtain variations using FreeBayes [4] version 0.9.21 from the short read alignments. We use vcffilter from vcflib [3] package version 1.0.2 to extract the SNPs We map the long reads using minimap2 [6] version 2.13-r858-dirty. Parameters for all tools are in 7. The short read and long read mapping, ploidy of the chromosome, and the SNPs are the inputs for HAT.

#### 2.1.2 Real data

We reconstruct the haplotypes of CBS1483, which is aneuploid and has ploidy ranging from one to five. It consists of ONT and paired-end Illumina reads that are available under the BioProject PRJNA522669. There are 4 ONT runs in this BioProject, and we used all of them in this study. Short reads have coverage of 159x and are 151bp, Nanopore reads have coverage of 72x with an average read length of 7kbp and N50 of 10kbp. We use the ASM1102231v1 assembly as the reference genome of CBS1483.

Moreover, we reconstruct the haplotypes of *Brettanomyces bruxellensis* strain GB54, a triploid genome which has higher heterozygosity, and longer chromosomes than CBS1483. The longest chromosome of GB54 is 4Mbp which is 3 times larger than CBS1483 largest chromosome. The ONT and paired-end Illumina reads are available under the BioProject PRJEB40511. The Illumina short reads are 75bp and 30x coverage. The nanopore long reads have the average read length of 12kbp, 82x coverage, and 23kbp N50. We use the DEBR_UMY321v1 assembly as the reference genome of GB54.

In both real datasets, the SNPs and the alignments are obtained with the same method as in the simulated datasets.

### 2.2 HAT method

HAT reconstructs haplotypes by linking alleles at SNP loci together using short and long reads. HAT comprises three components - initialization, iteration and assembly. Initialization creates the first phased blocks. The iteration expands the phased blocks and finds alleles of all haplotypes. Then, HAR clusters the reads, and assembles haplotypes using these clustered reads. An overview of the HAT algorithm can be seen in 1.

#### 2.2.1 Initialization

In initialization (see Fig. 1A), the multiplicity blocks are found, and then the first phased blocks are created. Phased blocks are a set of consecutive SNP loci in the phase matrix where the alleles are connected. First HAT creates seeds, which are a combination of consecutive SNP loci covered by the same short read; a single seed can be as small as two SNP loci. To create the seeds, we determine the SNP loci each short read is covering. If a read covers more than two SNP loci, we create all combinations of consecutive SNPs with different lengths and starting points. When we create a seed, based on the alleles present in the short reads that cover it, we obtain a set of combinations of alleles. While processing a short read, if the seed it would create already exists, no new seed is created and only the combination of alleles in the new read is added to the existing seed. In addition, we store the number of reads supporting each combination of alleles. Next, we filter the combination of alleles and the seeds. The combinations of alleles with fewer than five reads supporting are removed to remove erroneous combinations of alleles. If a seed ends up with fewer than two combinations of alleles, it is removed.

**Figure 1.**
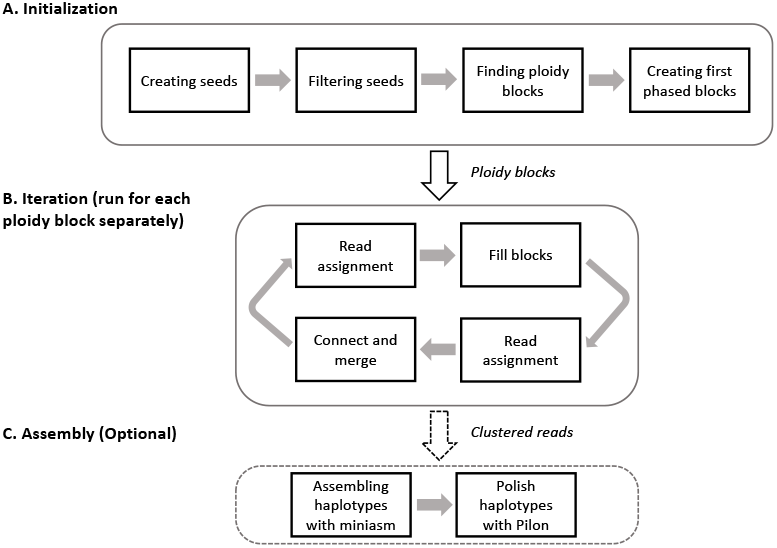
HAT overview. (A.) HAT creates seeds based on short read alignments and the location of SNPs. Then, it removes the combinations of alleles with low support as well as overlapping seeds. Next, HAT finds multiplicity blocks and creates the first phased blocks within them. (B.) HAT assigns reads to the blocks and haplotypes; based on these read assignments it fills the unphased SNPs within blocks. (C.) Finally, HAT can also use miniasm to assemble haplotype sequences for each block and polishes the assemblies using Pilon, but this step is optional

Next, HAT finds overlapping seeds and keeps only one of them because each SNP locus should be at most in one of the seeds to avoid conflicts. When HAT finds overlapping seeds, we check the number of combinations of alleles in each seed, the support of the seeds, and the first SNP locus of the seeds, then HAT picks the seed with the maximum number of combinations of alleles. If there are two overlapping seeds with the same number of combinations, we pick the longest one.

Then, HAT detects the regions that contain at least two different haplotypes, which we call multiplicity blocks. We use the sorted set of seeds as the input for Algorithm 1 to find multiplicity blocks and their corresponding ploidy. The seeds are sorted with respect to the number of combinations of alleles, and when two seeds have the same number, the one with an earlier position of the first SNP locus will come first. Once Algorithm 1 has identified the multi plicity blocks, if the estimated ploidy of the block exceeds the ploidy of the chromosome, it is decreased to the ploidy of the chromosome. In such cases, HAT eliminates combinations of alleles with low support until each seed has the same number of combinations as the chromosome’s ploidy. HAT creates a separate phase matrix for each multiplicity block. Each row of the phase matrix corresponds to one of the haplotypes, and each column is representative of an SNP locus within the multiplicity block.

Finally, HAT generates the first phased blocks. First, HAT removes the seeds with fewer combinations of alleles than the estimated ploidy of the block. Next, we use the combinations of alleles of the seeds to fill the phase matrix in the columns the seed is covering. Each seed creates a separate phased block because the relation of combinations of alleles of different seeds is unclear to one another. The SNP loci that do not belong to any block are added to the closest block to them.

#### 2.2.2 Iteration

The iterative part of HAT (Fig. 1B and Fig.2B) continues until there is only one block and the phase matrix is full, or if the blocks and phase matrix stop updating. We run the first iteration with short reads and the rest with long reads.

**Figure 2.**
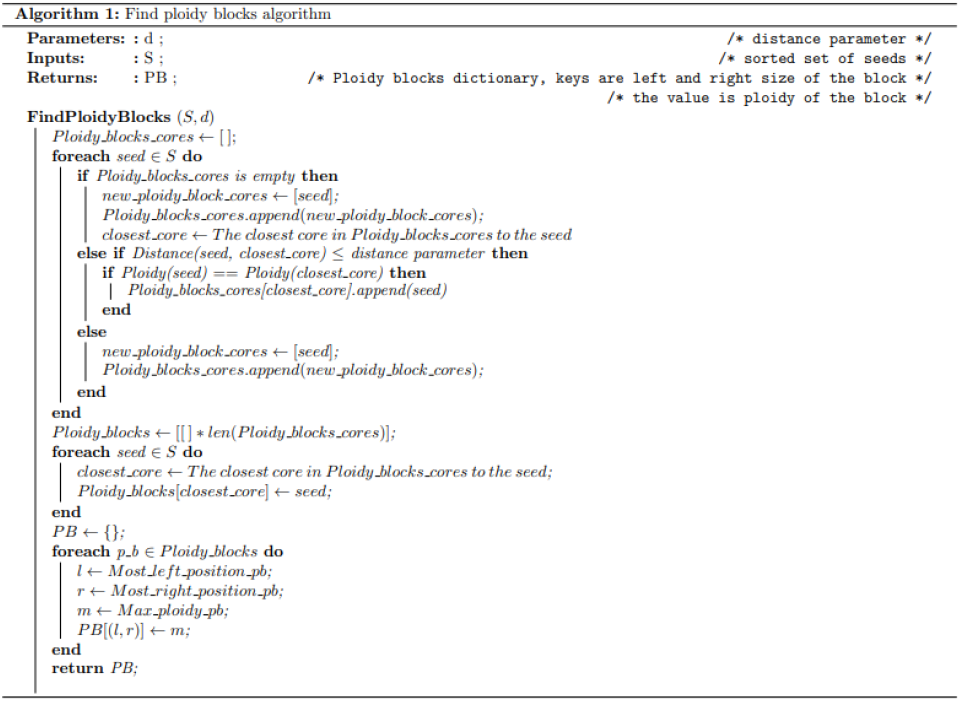
Find multiplicity blocks algorithm. The find multiplicity blocks algorithm takes the seeds and a distance parameter as input, and it returns multiplicity blocks as output. Each multiplicity block has a start and end position as well as an estimated ploidy for that region.

An essential step of the iterative HAT algorithm is assigning long and short reads to haplotypes in blocks. Each stage of the iterative part uses these assigned reads. Therefore, after each of the other main stages, reads are reassigned to the haplotype blocks based on the latest changes.

First, for every read we check the phased SNP loci that it covers within a block. If the combinations of alleles at those loci are unique for each haplotype, the read is assigned to the block. Then, the alleles of the read located at the phased SNP loci the read is covering within the block are compared with the alleles of each row of the phase matrix. The read is assigned to the haplotype if the Hamming distance to the row is less than distance_parameter, which changes with each run of the algorithm. When assigning long reads to the haplotypes, the distance_parameter is small (1) to accommodate sequencing errors. As the phasing algorithm proceeds, we increase distance_parameter to 3 to be stricter with the assignments, because there are more shared phased SNP locus within the block and the read.

The next stage of the iterative component is connecting and merging consecutive phased blocks. To connect two blocks, we use reads assigned to both blocks. We iterate over the blocks based on their location. If there is a one-to-one connection between all the haplotypes of two blocks with enough support, the blocks are merged, and the rows of the second block are switched so that the connected haplotypes of the first and second haplotypes are in the same row. Two haplotypes are connected if the number of reads supporting the connection is more than 1 in the first and second iteration, and 3 in the rest.

In the blocks’ filling step, we use all reads assigned to haplotypes of a block as input and process them to find the allele of unphased SNPs within the block by a majority voting between the reads of the haplotype. If the number of the reads supporting the majority vote allele is greater than 2, the allele is assigned to the haplotype’s SNP locus, and that cell of the phase matrix is filled. This phase might lead to some SNP loci being phased in some haplotypes but not in others. When iteration converges, HAT assigns long and short reads to haplotypes of each phased block using the read assignment module.

#### 2.2.3 Assembly

Optionally, HAT can assemble the reads to reconstruct sequence of the haplotypes using miniasm [12] version 0.3-r179 and the clustered long reads, then polish the assemblies using Pilon [13] version 1.24 and the clustered short reads. This part of HAT is optional. HAT uses miniasm and Pilon with default parameters. Users can use HAT to only cluster the reads and create the phase matrix and then use a tool of their choice to reconstruct sequence of the haplotypes.

### 2.3 Output

HAT outputs the following files:

- A multiplicity block figure which illustrates the multiplicity blocks and their level over the chromosome.
- The clustered reads files which contain the IDs of clustered reads for the haplotypes of each phased block.
- The phase matrix file which lists the alleles of haplotypes within each phased block.
- The haplotype sequences within each phased block. This output is optional, and it is produced only if the user also requests assembly.

### 2.4 Evaluating HAT

We run HAT version 0.1.7, nPhase version 1.1.10, and Whatshap polyphase version 0.19.dev161+g7660dcf on the simulated datasets and compare the phasing and read clustering accuracy. To calculate phasing accuracy of HAT, first we find a one-to-one mapping between the haplotypes HAT identifies within each block and the real haplotypes. Then we compare the allele of each haplotype at the SNP loci from the phase matrix to the ground truth. We count the number of correct SNPs for all the blocks and haplotypes and calculate accuracy as the count of correct SNPs divided by the total number of SNPs within the multiplicity blocks. To calculate the phasing accuracy of nPhase and Whatshap, we first find the haplotype which is the most similar to each cluster based on the cluster’s and haplotype’s alleles at the SNP loci the cluster covers. Then we devide the count of correctly phased SNPs by the total number of SNPs within the clusters.

Then, we assess the accuracy of read clustering. Since the simulated reads are already labeled with their native haplotype, we calculate the clustering accuracy by counting the number of reads clustered correctly. We also count the number of phased blocks to evaluate the reconstructed haplotypes’ completeness.

In addition to simulated data, we investigate the haplotypes HAT creates for the real CBS1483 and GB54 data.

## 3 Results

### 3.1 Conceptual overview of HAT using the example of a triploid chromosome

To provide an overview of the HAT algorithm, we consider chromosome ScII of triploid genome *Saccharomyces pastorianus* CBS1483 (ChrSc2), for a step-by-step discussion of HAT. The HAT algorithm consists of three main steps: (i) initialization, (ii) iteration and (iii) assembly. The input is a combination of both short and long reads, along with a reference genome. HAT will produce read clusters per haplotype when run in default settings. If the optional assembly parameter is supplied by the user HAT will also generate the haplotype sequences.

In the initialization step, HAT builds prototype phased blocks from seeds within multiplicity blocks. Phased blocks are fully resolved haplotype segments, while multiplicity blocks are genomic regions presenting sufficient variants for phasing and have an estimated ploidy associated. The initialization consists of three steps. First, HAT uses the alignment of short reads to the reference to find well-supported combinations of variant alleles, called seeds. In our example of ChrSc2, 528 SNPs were used to create 25335 combinations, which are filtered down to 119 by removing the combinations with low support (see Methods). Next, HAT constructs multiplicity blocks from the seeds and their corresponding ploidy with Algorithm 1. Finally, HAT uses seeds with matching number of combinations of alleles to create the first phased blocks within each multiplicity block. In the example of ChrSc2, HAT found 16 multiplicity blocks (Fig. 4).

During the iterative phase HAT processes each multiplicity block to phase the remaining SNPs and create bigger phased blocks within a multiplicity block. It consists of two sections: (i) filling blocks, and (ii) merging blocks. Before running each section, HAT assigns reads to blocks and haplotypes based on the SNPs each read covers and their similarity to the phased SNPs. 8 shows how each step of the iterative algorithm improves the phasing of ChrSc2. The iteration stops when there is no improvement over the previous step. The first iteration uses the short reads, while the remaining iterations use long reads. In our experiments with real and simulated data, HAT converges in less than four iterations. Increasing the number of iterations for the short reads does not change the overall phasing performance, because the blocks are bigger than the linking range of the short reads.

Upon convergence, there are 23 phased blocks and only 23 unphased alleles from the SNP loci within the multiplicity blocks. We use miniasm on the long reads assigned to haplotypes of each phased block to assemble them. Then, we polish the assemblies with the short reads assigned to haplotypes using Pilon.

### 3.2 HAT outperforms state-of-the-art on simulated data

To evaluate HAT we use simulated datasets, consisting of short and long reads, and alignments to the haplotypes. Details of simulation are described in the Methods section. Summary statistics of the simulated data sets are reviewed in Table 2.

We compare HAT to nPhase and Whatshap polyphase using the various metrics (see Methods); Table 3 summarizes the performance of the tools on the simulated data sets. First, we compare long read clustering accuracy. The number of long reads clustered incorrectly by HAT is lower than that of both nPhase and Whatsap for all ploidy levels: HAT’s error rate ranges from less than 1% (triploid high heterozygosity) to 24% (pentaploid low heterozygosity), whereas for nPhase the range is 5% (tetraploid high heterozygosity) to 38% (pentaploid low heterozygosity) and for Whatshap it is 7% (tetraploid high heterozygosity) to 22% (triploid high heterozygosity).

For all datasets, HAT successfully phases at least 90% of the SNPs, and the accuracy is the highest at 98% for the triploid high heterozygous genome (last column in Table 3). In all datasets, Whatshap has the lowest accuracy and HAT has the highest. Note that the phasing accuracy of HAT is calculated only for the SNPs inside the multiplicity blocks, but the multiplicity blocks cover almost all of the SNPs on the chromosome, with the lowest coverage being 78% for the pentaploid low heterozygous genome (Table 1). Similarly, for nPhase and Whatshap, we calculated the phasing accuracy based on the clusters each tool generates.

**Table 1.** The percentage of SNPs that are inside multiplicity blocks

| Dataset | Percentage of SNPs inside multiplicity blocks |
| --- | --- |
| Triploid low heterozygosity | 92% |
| Triploid high heterozygosity | 99% |
| Tetraploid low heterozygosity | 87% |
| Tetraploid high heterozygosity | 99% |
| Pentaploid low heterozygosity | 78% |
| Pentaploid high heterozygosity | 98% |

**Table 2.** Descriptive statistics of the simulated datasets and ChrSc2, the base chromosome used for simulations.

| Dataset | Ploidy | Simulated SNPs | # SNPs found by FreeBayes | # short reads | # long reads |
| --- | --- | --- | --- | --- | --- |
| Triploid low heterozygosity | 3 | 1230 | 687 | 194910 | 3295 |
| Triploid high heterozygosity | 3 | 6398 | 4143 | 194910 | 3441 |
| Tetraploid low heterozygosity | 4 | 1144 | 506 | 259880 | 4423 |
| Tetraploid high heterozygosity | 4 | 12072 | 7512 | 259880 | 4358 |
| Pentaploid low heterozygosity | 5 | 1606 | 504 | 324850 | 5433 |
| Pentaploid high heterozygosity | 5 | 17802 | 7232 | 324850 | 5394 |
| CBS1483 chromosome ScII | 3 | — | 528 | 428802 | 8051 |

**Table 3.** HAT outperforms nPhase and Whatshap in phasing accuracy on simulated data. The first two columns show number reads clustered and phased, third column lists the number of SNPs within multiplicity blocks that were phased correctly/incorrectly and unphased. The final column is the SNP phasing accuracy.

| Dataset | Tool | Read clustering error |  | Total reads phased |  | SNP phasing performance |  |  | Accuracy<sup>2</sup> |
| --- | --- | --- | --- | --- | --- | --- | --- | --- | --- |
|  |  | short<sup>1</sup> | long | short<sup>1</sup> | long | correct | incorrect | unphased |  |
| Triploid low | HAT | 14 | 65 | 5400 | 2122 | 1813 | 20 | 35 | 98% |
|  | nPhase | – | 307 | – | 1268 | 1936 | 379 | 0 | 84% |
|  | Whatshap | – | 225 | – | 1943 | 759 | 1215 | 0 | 61% |
| Triploid high | HAT | 55 | 13 | 37291 | 2138 | 11895 | 185 | 19 | 98% |
|  | nPhase | – | 218 | – | 2829 | 12701 | 1464 | 0 | 90% |
|  | Whatshap | – | 680 | – | 3060 | 7106 | 4663 | 0 | 60% |
| Tetraploid low | HAT | 135 | 187 | 2466 | 1580 | 1549 | 34 | 68 | 94% |
|  | nPhase | – | 297 | – | 1439 | 2023 | 335 | 0 | 86% |
|  | Whatshap | – | 254 | – | 2229 | 1434 | 538 | 0 | 72% |
| Tetraploid high | HAT | 62 | 32 | 17968 | 2252 | 29039 | 493 | 37 | 98% |
|  | nPhase | – | 219 | – | 4053 | 28980 | 1990 | 0 | 94% |
|  | Whatshap | – | 287 | – | 4142 | 20550 | 7942 | 0 | 72% |
| Pentaploid low | HAT | 545 | 518 | 2350 | 2141 | 1341 | 51 | 74 | 91% |
|  | nPhase | – | 662 | – | 1726 | 1714 | 287 | 0 | 86% |
|  | Whatshap | – | 348 | – | 2396 | 1803 | 547 | 0 | 76% |
| Pentaploid high | HAT | 1041 | 266 | 9360 | 6804 | 31980 | 1479 | 224 | 95% |
|  | nPhase | – | 449 | – | 5264 | 35450 | 2676 | 0 | 93% |
|  | Whatshap | – | 708 | – | 5246 | 27929 | 7301 | 0 | 79% |

<sup>1</sup> The short read cluster error was calculated only for HAT because nPhase and whatshap are designed specifically to cluster the long reads.
<sup>2</sup> Accuracy is defined in Methods.

To assess the phasing contiguity we checked the number of phased blocks in the HAT output, and report that for highly heterozygous cases HAT can phase almost all of the haplotypes. HAT creates 2 phased blocks for the triploid case and 3 phased blocks for the tetraploid one. For cases with low heterozygosity, HAT creates 15, 30 and 23 phased blocks for the triploid, tetraploid and pentaploid genomes. This is expected because these genomes are largely identical and it is not possible to connect the phased blocks. In contrast, we also note that for the highly heterozygous pentaploid dataset, HAT creates 33 phased blocks although it has 96% phasing accuracy, an outcome likely caused by the high ploidy level.

### 3.3 HAT shows robust performance on real data

Since there are not many chromosome-level polyploid assemblies available, the disparity between simulated and real genomes can be significant. Hence, we are evaluating HAT on the real *Saccharomyces pastorianus* CBS1483 dataset to corroborate the results from the simulated datasets as well. CBS1483 is a valid test model because it is aneuploid and has various ploidies ranging from one to five. Additionally, the chromosomes are small and easy to investigate. We report read clustering and phasing results for seven chromosomes of CBS1483 representing various levels of ploidy, heterozygosity and length in Table 4. For the highly heterozygous chromosome ScI, multiplicity blocks that HAT finds cover 54% of the whole sequence and within these blocks HAT phased 96% of the alleles. Although ScIV contains a large number of SNPs, it is the largest chromosome and all the SNPS are concentrated around the centromere and thus, the % of phased regions is lower. SeI, on the other hand, is one of the shortest chromosomes (185kb long) and there are very few SNPs, meaning that the haplotypes are identical in most positions on the chromosome. For that reason, HAT phases less than 1% of the chromosome.

**Table 4.** HAT results on CBS1483 real data. These chromosomes are a representative subset of all chromosomes of CBS1483

| Chromosome name | Ploidy | # SNPs | % phased regions | Alleles within blocks |  |
| --- | --- | --- | --- | --- | --- |
|  |  |  |  | Phased | unphased |
| ScI | 3 | 619 | 54% | 1384 | 60 |
| ScII | 3 | 528 | 14% | 922 | 23 |
| ScIV | 3 | 1643 | 20% | 3424 | 168 |
| ScIX | 2 | 195 | 10% | 335 | 43 |
| ScVIII | 5 | 417 | 14% | 687 | 4 |
| SeI | 2 | 21 | <1% | 8 | 0 |
| SeVII-ScVII | 3 | 341 | 4% | 444 | 5 |

We observe the haplotype sequences created by miniasm and polished by Pilon for the multiplicity block 153738,163604 in chromosome ScII (Fig. 4). This multiplicity block is only 8kb long, and the estimated ploidy for that region is 2. To visually investigate the accuracy of haplotype reconstruction, we map the clustered reads to haplotype 1 and haplotype 2 reconstructed by HAT and view the alignment in Integrative Genome Viewer (iGV). Fig. 4 depicts the alignment of clustered reads of CBS1483 Chromosome ScII to the sequence of the first haplotype. The reads that belong to each haplotype have matching alleles that can differentiate them from reads of other haplotypes. We, therefore, demonstrate that the HAT algorithm for read clustering and finding multiplicity blocks works on real data.

Previous studies shows that the UIP3 gene is removed in some of the haplotypes of CBS1483 chromosome ScI [10]. We investigate the same gene in HAT output by aligning the reads HAT clustered in the multiplicity block covering the positions from 169765 to 178549 where UIP3 is located (see 7), to UIP3 sequence. As expected, only haplotype 2 reads align to the gene.

Finally, we test the performance of HAT on GB54, a triploid *Brettanomyces bruxellensis* yeast strain, which Abou Saada et al. also uses to evaluate nPhase [1]. GB54 is an interesting test set because it has longer chromosomes and the chromosomes are more heterozygous compared to CBS1483. Table 5 shows HAT’s performance in phasing GB54, and 8 illustrates the multiplicity blocks HAT finds. As expected, the percentage of phased regions is much larger than that of CBS1483 (Table 4), since GB54 is more heterozygous. Additionally, when we visualize the multiplicity blocks in GB54 we observe multiple long regions in chromosome 2, 3, and 5 where two of the haplotypes are identical. For instance, on Chr 3 the genomic region from 909325 to 1172321, all of the seeds have only two combination of alleles, and the average ratio of the read support for combination of alleles of seeds at these regions is 1.7. This is in line with our expectation that two of the haplotypes are identical in this region, and we get on average near twice as much read coverage for one of the haplotypes as there would be if there were three different haplotypes. Morever, when Abou Saada et al. phased Chr 4 using nPhase they reported that two of the haplotypes were identical at the end region of Chr 4 since they could phase only two haplotypes. However, when we look at the same location on Chr 4, we observe two small genomic regions (from 1407542 to 1430976 and from 1533678 to 1546306) where HAT can successfully phase all three haplotypes (see 8).

**Table 5.** HAT results on GB54 real data.

| Chromosome | # SNP loci | % of phased regions | Alleles within blocks |  |
| --- | --- | --- | --- | --- |
|  |  |  | Phased | Unphased |
| Chr 1 | 36628 | 84% | 96764 | 2001 |
| Chr 2 | 23756 | 82% | 60066 | 2586 |
| Chr 3 | 18902 | 86% | 35728 | 1807 |
| Chr 4 | 19377 | 89% | 50616 | 884 |
| Chr 5 | 10609 | 63% | 17867 | 1673 |
| Chr 6 | 15762 | 72% | 32877 | 1295 |
| Chr 7 | 3581 | 92% | 10482 | 216 |
| Chr 8 | 1327 | 72% | 2949 | 305 |

## 4 Conclusion

HAT is a haplotype assembly tool that reconstructs haplotypes and phases genomes using NGS and TGS data. It is impossible to phase entire homologous chromosomes when there are large variation deserts. To address this, HAT identifies regions where some of the haplotypes are identical so they are taken into account when phasing. We show that NGS and TGS provide enough information to phase high heterozygosity genomes on a chromosome-scale and more than 90% of the alleles in a low heterozygosity genomes.

We evaluate the performance of HAT on six simulated datasets based on an aneuploid yeast strain *Saccharomyces pastorianus* CBS1483, and compare it to nPhase and Whatshap, the state-of-the-art algorithms. We observe that HAT presents higher phasing accuracy, which results from starting with seeds created by accurate short reads. While all tools have decent performance in highly heterozygous genomes, HAT performs remarkably well in phasing and read clustering of low heterozygote genomes. However, in the latter case, haplotypes created by HAT are fragmented since it does not attempt to connect the multiplicity blocks, because there is not enough information to link them. While we did not evaluate HAT on any diploid dataset directly, we observed that HAT successfully phases blocks with multiplicity level of 2 which shows that it can also be applied to diploid genomes.

The main limitation of HAT is that it uses of alignments of short and long reads to the reference. Similar to other haplotype assembly tools, HAT’s performance greatly depends on the quality of this alignment and the subsequent variant calling. Moreover, HAT uses only the SNPs for phasing, thus it may not be able to reconstruct haplotypes in genomes with high levels of insertions, deletions, and structural variations.

Another potential limitation is that in rare cases, HAT may incorrectly assign a lower than actual multiplicity number. This occurs when there is a group of seeds in close proximity where the number of combinations of alleles in any of the seeds is smaller than the actual multiplicity level. When each seed is viewed separately, some of the haplotypes appear to be identical in that region. However, it is possible that these are different groups of identical haplotypes, and the ploidy level of the region may be higher if all of these seeds are viewed as a whole. This can be solved by creating seeds and identifying multiplicity blocks using long reads. However, considering all consecutive SNP loci in the reads as seeds requires significant computing power since each long read might cover hundreds of SNPs. Additionally, allelic combinations in the seeds may be affected by the high error rate of long reads. Another way to mitigate erroneous multiplicity assignment is to adjust multiplicity levels during the iterative part of HAT when long reads are used to phase the SNPs. In principle, by solving the mentioned problem it should be possible to create the seeds with HiFi reads, which will lead to longer multiplicity blocks and higher contiguity.

There are not many polyploid haplotype resolved genomes at the chromosome scale, which hinders the development of novel haplotype assembly algorithms. Hence, haplotype simulators are limited and the simulated haplotypes differ significantly from the real ones. We observed this when we compared the multiplicity blocks of real and simulated data (compare Fig. 3 and 6). There are many regions in the real data where the multiplicity level is smaller than the chromosome’s actual ploidy level, contrary to simulated data. This might change with HiC reads since they provide long range information and link regions of the chromosome that are far apart. In addition to inconsistencies in the ploidy levels, the large variation deserts in CBS1483 genome cannot be simulated due to limitations of current haplotype simulators.

**Figure 3.**
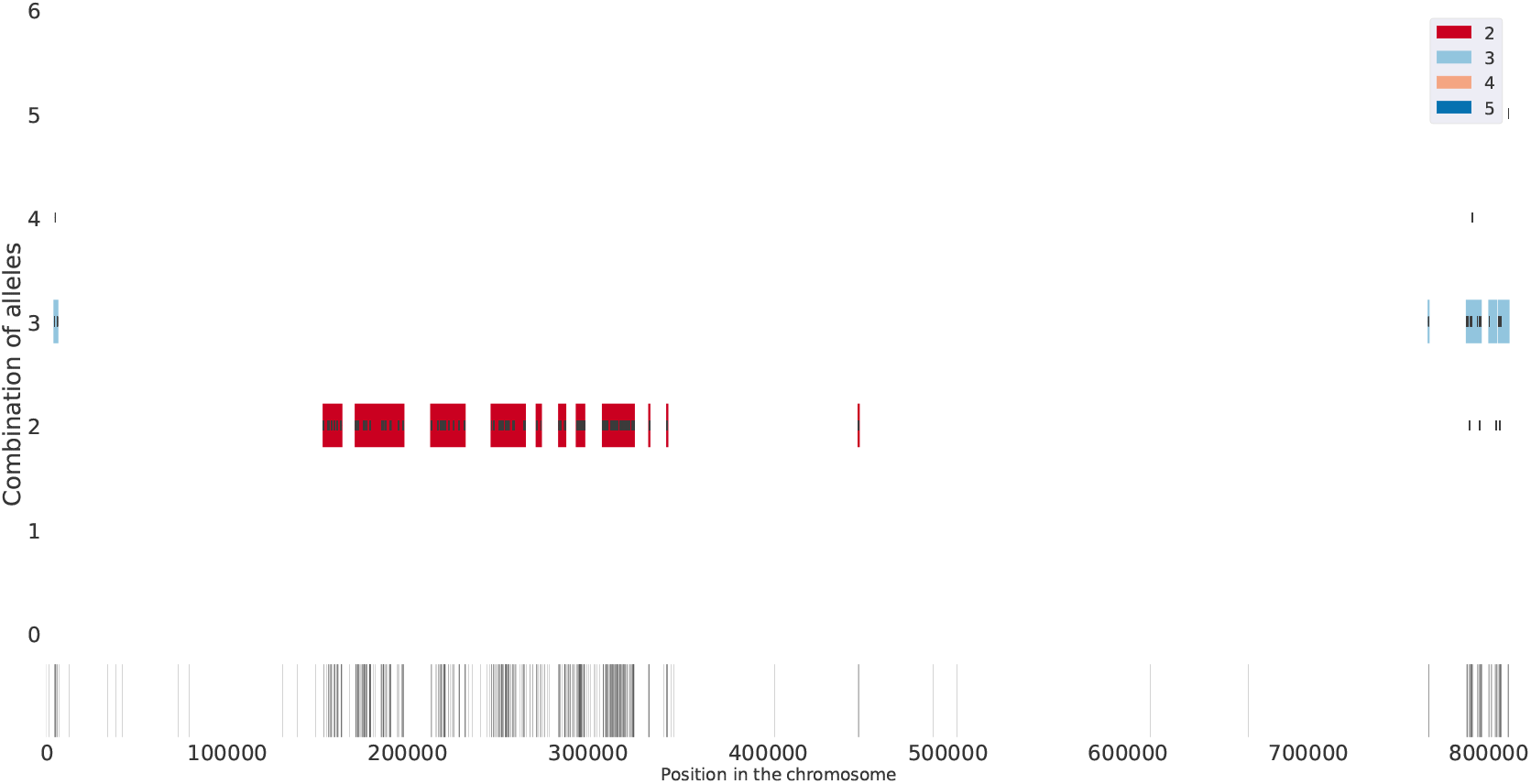
Multiplicity blocks HAT finds for ChrSc2 of CBS1483. The output of finding multiplicity blocks algorithm on real data, ChrSc2 of CBS1483. The long, black vertical lines at the bottom show the SNPs and their positions on the chromosome found by FreeBayes. From these SNPS, HAT finds the seeds shown in short, black vertical lines in panel above the SNPs. The seeds are placed vertically based on the number of combination of alleles they have, ranging from 1 to 6 (y axis). HAT uses these seeds to find multiplicity blocks, which are shaded regions encapsulating the seeds and the color of the region indicates the estimated ploidy level. See the legend for the colors corresponding to different ploidy levels.

**Figure 4.**
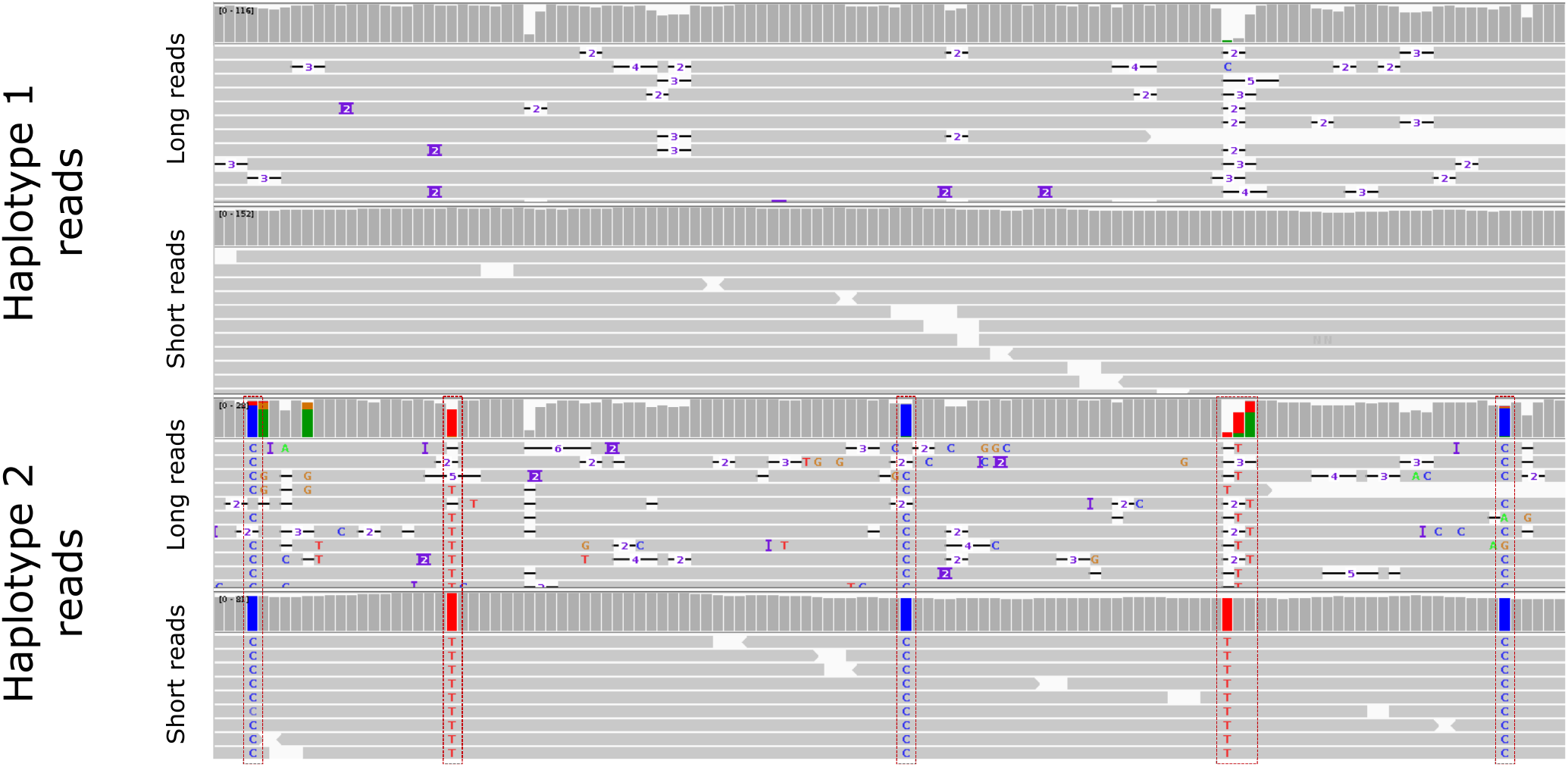
HAT can accurately cluster reads to reconstructed haplotypes. We aligned short and long reads to haplotype 1 (top two rows) and haplotype 2 (bottom two rows) phased by HAT for the multiplicity block covering the positions from 153738 to 163604 of ChrSc2 and visualized the alignment using iGV. Haplotype 2 reads differs significantly from haplotype 1 reads at five positions (outlined with red rectangles).

Although we demonstrate the performance of HAT on only two yeast strains *Saccharomyces pastorianus* CBS1483 and *Brettanomyces bruxellensis* GB54, HAT can also phase different polyploid genomes. Since HAT performed consistently well on various levels of ploidy and heterozygosity, we expect our results to generalize to other genomes of varying ploidy and that HAT can readily be adopted to different use-cases. Moreover, we presume that HAT can find applications in metagenomics assembly since the haplotype and metagenomics assembly problems are comparable at the strain level. In metagenomics assembly, the goal is to reconstruct the genome of every single strain of the metagenomics community, which can be up to thousands of genomes. These strains, like haplotypes, are quite similar to each other. Furthermore, as a result of horizontal gene transfers, some of the species within the community share genomic content, complicating their read separation. Moreover, the sequencing coverage of strains varies significantly, which might lead to the underrepresented strains not being reconstructed in the assembly process.

Ultimately, HAT enables us to reconstruct haplotypes of polyploid genomes reliably, investigate the relationship of phenotypic features to the underlying haplotype alleles, and gain a better understanding of genetic diversity.

## Acknowledgements

We thank Erin Jordan and Stephanie Pillay for reviewing the paper.

## 5 Supplementary materials

**Table 6.**
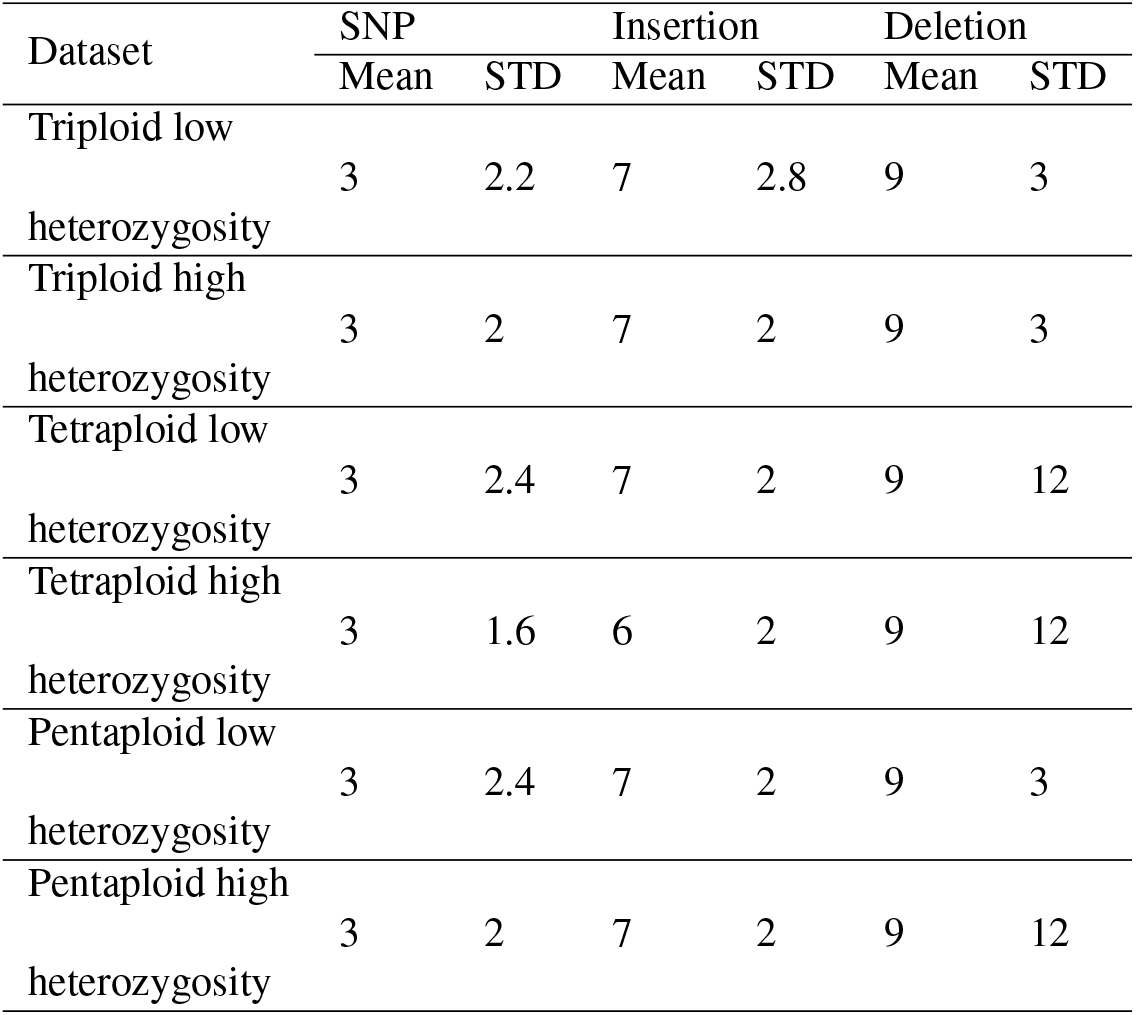
Haplogenerator parameters for simulating datasets.

**Table 7.** Parameters of the tools we use in this study.

| Tool name | Parameter name | Parameter value |
| --- | --- | --- |
| ART | Read length | 125 |
|  | Mean insertion size | 400 |
|  | Standard deviation of insertion size | 20 |
|  | coverage | 20 |
| Badread | quantity | 20 |
| Minimap2 | Secondary | No |
| BWA mem | Default | - |
| Vcfilter | -f | TYPE = SNP |
| FreeBayes | Ploidy | Ploidy of the dataset |
|  | Mean alternate count | 5 |
| Miniasm | Default | - |
| Pilon | Default | - |

**Table 8.**
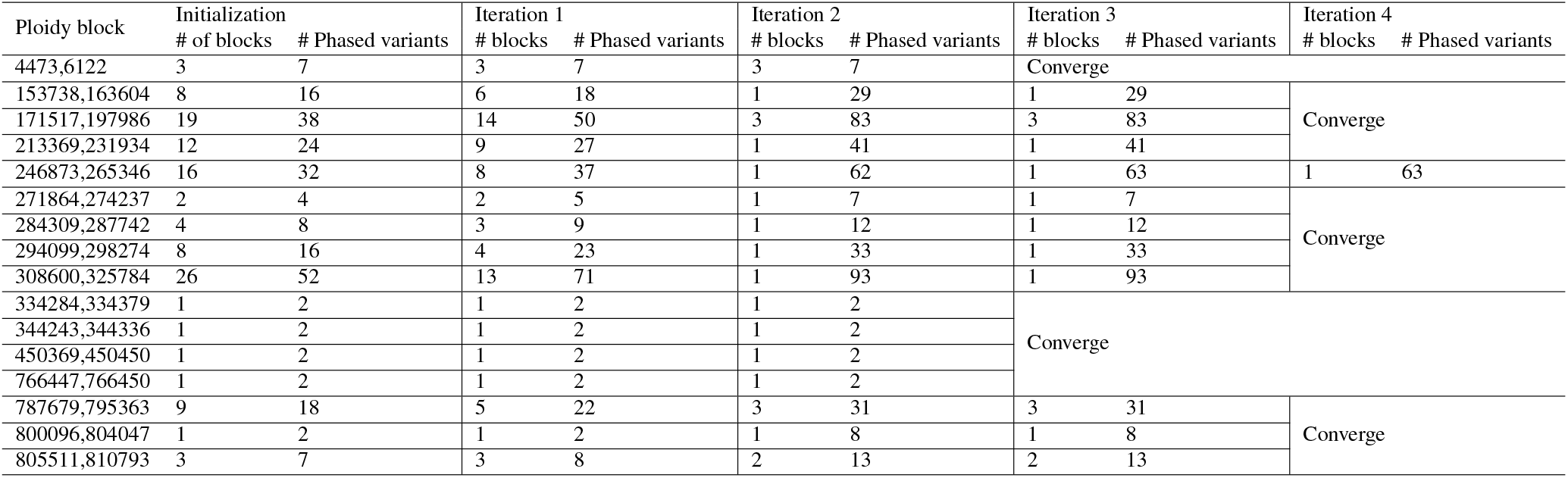
The haplotype reconstruction improvement after each step: we ran HAT on Chromosome ScII of CBS1483 and calculated the number of blocks and number of variants after each iteration of the iterative part of HAT. For all ploidy blocks HAT converges in at most 4 iterations.

**Figure 5.**
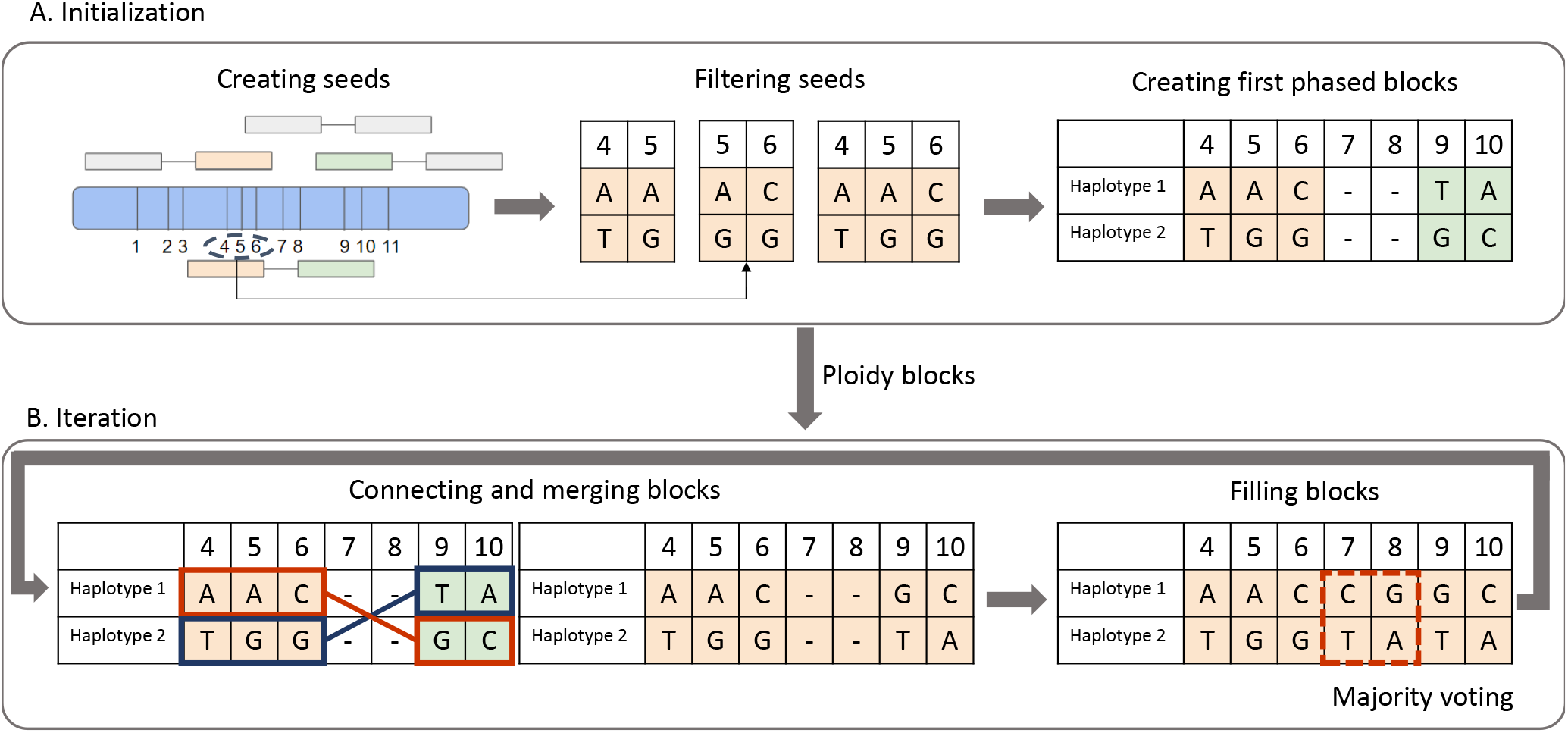
Iteration example. A. Based on the alignment of the reads that are covering SNPs 4,5 and 6, HAT creates three seeds. In this scenario, because the support of the combinations of alleles of these three seeds are equal, HAT keep the longer one in the filtering seeds step. After that, HAT use the remaining seeds and the combinations of alleles to create first phased blocks. B. Based on the read assignment, the reads that belong to haplotype 1 of block 1, also belong to haplotype 2 of block 2. This means, these two haplotypes are the same, and in the connecting and merging blocks, a bigger block is created, and the mentioned haplotypes are linked together. After that, based on the assignment of reads to each haplotype, and a majority voting between those reads, HAT finds the allele of the unphased SNPs.

**Figure 6.**
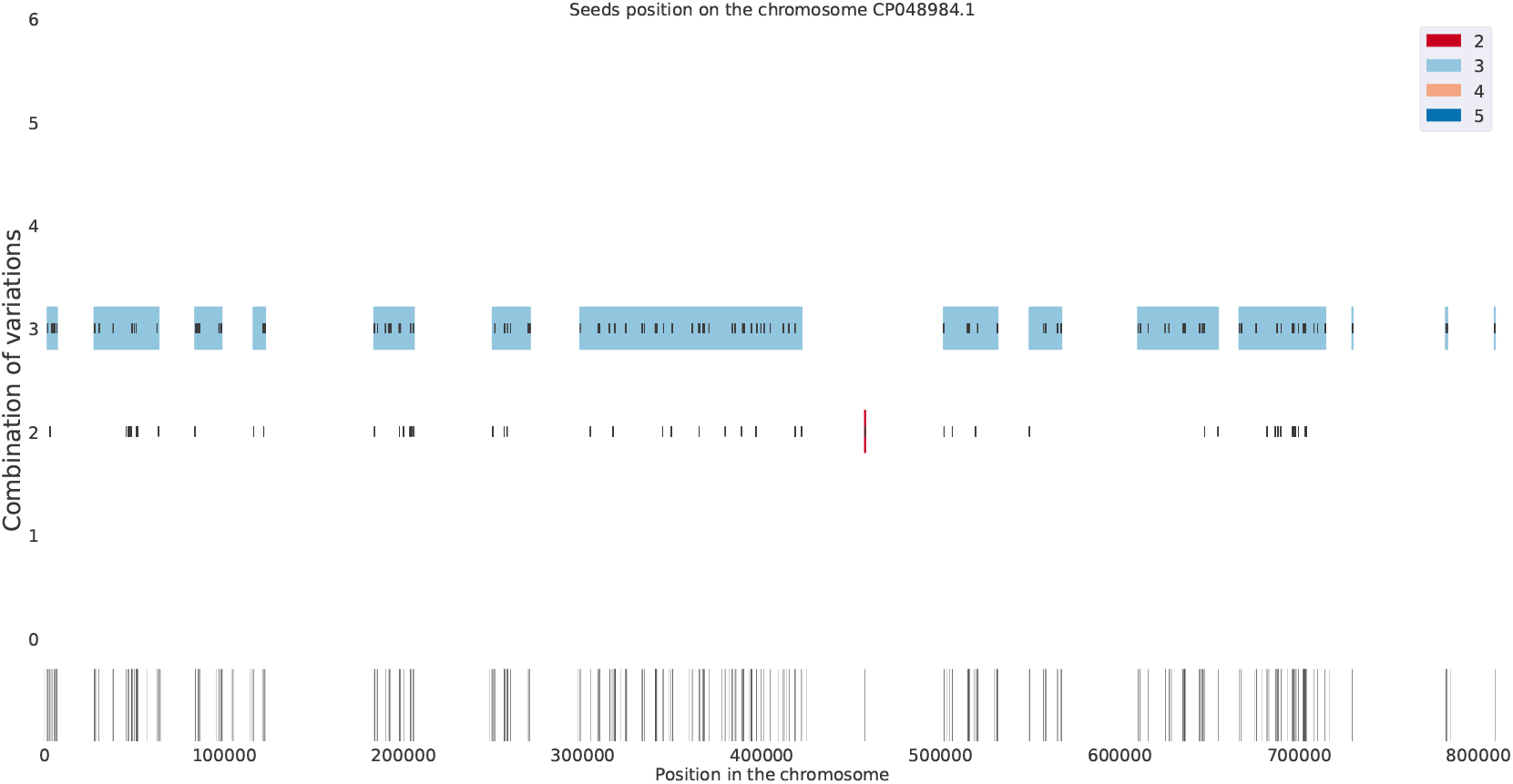
Multiplicity blocks of Triploid low heterozygosity dataset.

**Figure 7.**
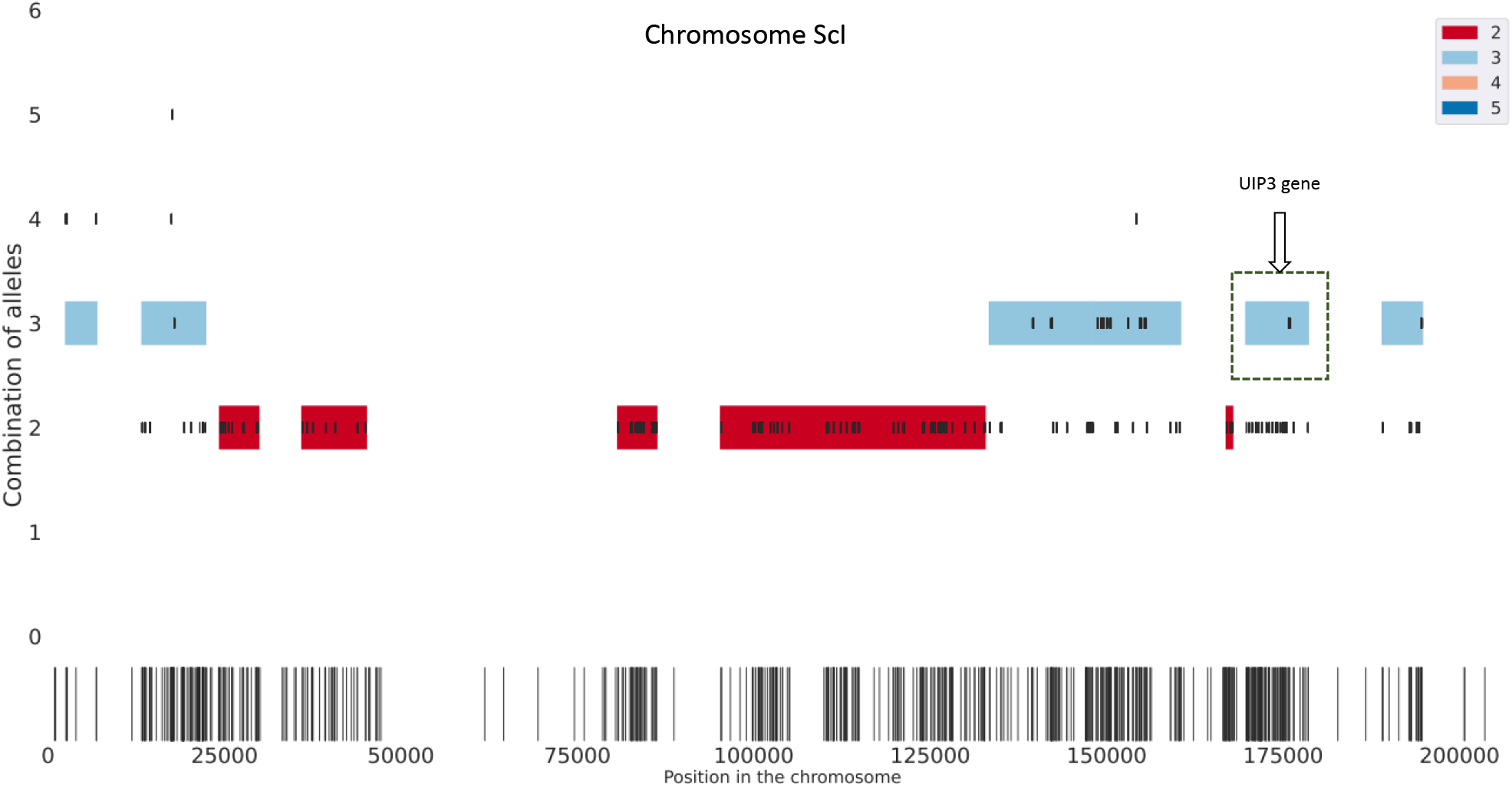
Multiplicity blocks of triploid ScI of CBS1483. Inside the green box you can find the multiplicity block which contains UIP3 gene.

**Figure 8.**
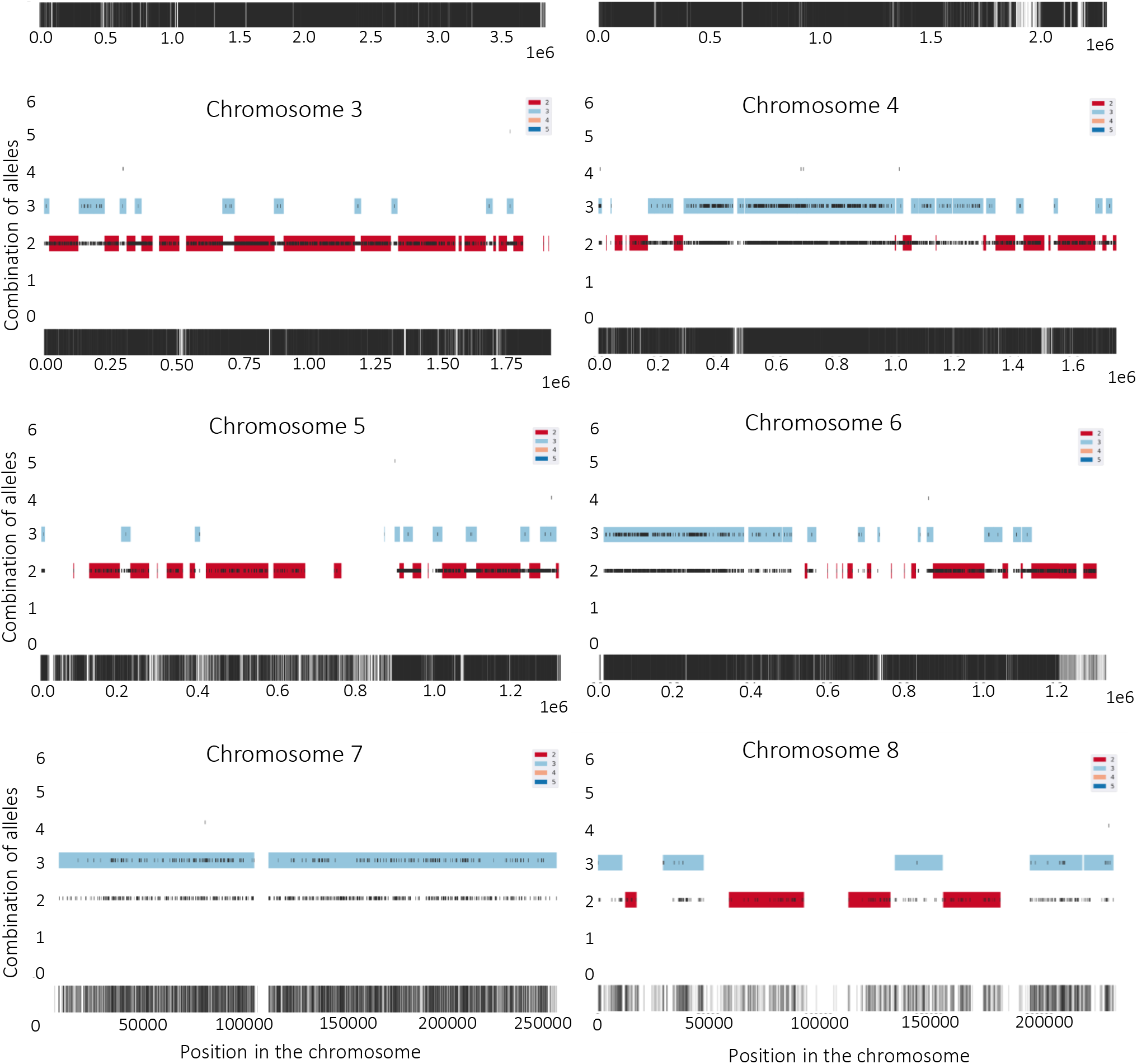
Multiplicity blocks of all chromosomes of triploid GB54.

## References

[1] O. Abou Saada, A. Tsouris, C. Eberlein, A. Friedrich, and J. Schacherer. nPhase: an accurate and contiguous phasing method for polyploids. Genome Biology, 22(1):126, Apr. 2021.

[2] S. Garg. Computational methods for chromosome-scale haplotype reconstruction. Genome biology, 22(1):1–24, 2021.

[3] E. Garrison, Z. N. Kronenberg, E. T. Dawson, B. S. Pedersen, and P. Prins. Vcflib and tools for processing the VCF variant call format. Technical report, bioRxiv, May 2021. Section: New Results Type: article.

[4] E. Garrison and G. Marth. Haplotype-based variant detection from short-read sequencing. arXiv:1207.3907 [q-bio], July 2012. arXiv: 1207.3907.

[5] W. Huang, L. Li, J. R. Myers, and G. T. Marth. ART: a next-generation sequencing read simulator. Bioinformatics, 28(4):593–594, Feb. 2012.

[6] H. Li. Minimap2: pairwise alignment for nucleotide sequences. Bioinformatics, 34(18):3094–3100, Sept. 2018.

[7] M.-H. Moeinzadeh, J. Yang, E. Muzychenko, G. Gallone, D. Heller, K. Reinert, S. Haas, and M. Vingron. Ranbow: A fast and accurate method for polyploid haplotype reconstruction. PLOS Computational Biology, 16(5):e1007843, May 2020. Publisher: Public Library of Science.

[8] E. Motazedi, R. Finkers, C. Maliepaard, and D. de Ridder. Exploiting next-generation sequencing to solve the haplotyping puzzle in polyploids: a simulation study. Briefings in Bioinformatics, 19(3):387–403, May 2018.

[9] J. Ramsey and D. W. Schemske. Pathways, Mechanisms, and Rates of Polyploid Formation in Flowering Plants. Annual Review of Ecology and Systematics, 29(1):467–501, 1998. _eprint: https://doi.org/10.1146/annurev.ecolsys.29.1.467.

[10] A. N. Salazar, A. R. Gorter de Vries, M. Van Den Broek, N. Brouwers, P. de la Torre Cortès, N. G. Kuijpers, J.-M. G. Daran, and T. Abeel. Chromosome level assembly and comparative genome analysis confirm lager-brewing yeasts originated from a single hybridization. BMC genomics, 20(1):1–18, 2019.

[11] S. D. Schrinner, R. S. Mari, J. Ebler, M. Rautiainen, L. Seillier, J. J. Reimer, B. Usadel, T. Marschall, and G. W. Klau. Haplotype threading: accurate polyploid phasing from long reads. Genome biology, 21(1):1–22, 2020.

[12] R. Vaser, I. Sović, N. Nagarajan, and M. Šikić. Fast and accurate de novo genome assembly from long uncorrected reads. Genome Res., 27(5):737–746, May 2017. Company: Cold Spring Harbor Laboratory Press Distributor: Cold Spring Harbor Laboratory Press Institution: Cold Spring Harbor Laboratory Press Label: Cold Spring Harbor Laboratory Press Publisher: Cold Spring Harbor Lab.

[13] B. J. Walker, T. Abeel, T. Shea, M. Priest, A. Abouelliel, S. Sakthikumar, C. A. Cuomo, Q. Zeng, J. Wortman, S. K. Young, and A. M. Earl. Pilon: An Integrated Tool for Comprehensive Microbial Variant Detection and Genome Assembly Improvement. PLoS ONE, 9(11):e112963, Nov. 2014.

[14] R. Wick. Badread: simulation of error-prone long reads. JOSS, 4(36):1316, Apr. 2019.

